# Avoiding ascertainment bias in the maximum likelihood inference of phylogenies based on truncated data

**DOI:** 10.1101/186478

**Authors:** Asif Tamuri, Nick Goldman

## Abstract

Some phylogenetic datasets omit data matrix positions at which all taxa share the same state. For sequence data this may be because of a focus on single nucleotide polymorphisms (SNPs) or the use of a technique such as restriction site-associated DNA sequencing (RADseq) that concentrates attention onto regions of differences. With morphological data, it is common to omit states that show no variation across the data studied. It is already known that failing to correct for the ascertainment bias of omitting constant positions can lead to overestimates of evolutionary divergence, as the lack of constant sites is explained as high divergence rather than as a deliberate data selection technique. Previous approaches to using corrections to the likelihood function in order to avoid ascertainment bias have either required knowledge of the omitted positions, or have modified the likelihood function to reflect the omitted data. In this paper we indicate that the technique used to date for this latter approach is a conditional maximum likelihood (CML) method. An alternative approach — unconditional maximum likelihood (UML) — is also possible. We investigate the performance of CML and UML and find them to have almost identical performance in the phylogenetic SNP dataset context. We also make some observations about the nucleotide frequencies observed in SNP datasets, indicating that these can differ systematically from the overall equilibrium base frequencies of the substitution process. This suggests that model parameters representing base frequencies should be estimated by maximum likelihood, and not by empirical (counting) methods.

## Introduction

Leaché et al. (2015) considered likelihood methods that are available for the phylogenetic analysis of single nucleotide polymorphism (SNP) data sets, i.e. nucleotide (nt) sequence alignments in which any constant sites have been omitted. (Note that the term ‘constant site’ is used to mean one in which no differences are apparent amongst the sequences collected, and not to suggest that substitution cannot ever occur there.) Starting with an equivalent situation in the analysis of restriction sites (Felsenstein, 1992), then with SNP data (Kuhner et al., 2000), morphological data (Lewis, 2001) and most recently with restriction site-associated DNA sequencing (RADseq; Baird et al., 2008; Seeb et al., 2011; Peterson et al., 2012), it has been clear that omitting such sites is a form of ascertainment bias. Analyzing only variable sites, without correction, can lead to overestimation of branch lengths and biases in phylogeny inference (Lewis, 2001; Leaché et al., 2015).

Leaché et al. (2015) investigated three likelihood methods for correcting for the omission of constant sites. The first, developed by Felsenstein (1992) and Lewis (2001) and denoted lewis by Leaché et al. (2015), uses a conditional likelihood and does not explicitly consider the number of constant sites missing from the data set. The second, described by Kuhner et al. (2000) and denoted felsenstein by Leaché et al. (2015), uses a ‘reconstituted likelihood’ requiring the total number of constant sites to be known, but does not consider their partitioning into constant-A,-C,-G or-T sites. The third method (stamatakis: Leaché et al., 2015) again uses reconstituted likelihood, now requiring knowledge of the exact numbers of each of the four different constant site patterns. The result of this approach is necessarily exactly the same as analyzing the original data set, with no sites omitted.

The felsenstein and stamatakis methods can be used in cases where data (constant sites) are omitted but the numbers of such sites are known — situations that are unlikely to arise with modern data recording techniques. Situations with unknown amounts of omitted data are more frequent, and warrant further attention. In this paper we place the lewis method into a more-general likelihood inferential framework and derive a new method for estimating parameters (e.g. tree topologies, branch lengths and substitution model parameters), as well as the number of omitted sites in the case that this is not known. The new method performs almost identically to the lewis method, and we explore the reasons for this. Lastly, we make some observations about the effect that SNP data (i.e. missing observations of constant sites) have on observed base frequencies.

## Methods

Using a slightly different notation from Leaché et al. (2015) to describe the inference problem, we assume a total of *n* observations (SNP or non-constant sites), of which *n*_*i*_ correspond to pattern (possible alignment column) *i*; if we observe *l* different patterns, then 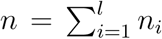. (Sums and products are generally over *i* = 1,…, *l* throughout this paper; for simplicity we omit these limits when there is no ambiguity) For a SNP data set, we do not observe the constant-A,-C,-G or-T patterns; we write the unobserved number of these as *m*, and the total number of sites (observed and unobserved) as *n* + *m* = *N*. Such data sets are described as *truncated* (Blumenthal, 1981).

To complete the description, we need a model describing the probabilities of occurrence of both the observed and unobserved patterns. As usual in likelihood-based phylogenetics, we will assume that an underlying tree structure with branch lengths is to be estimated, along with any free parameters of a substitution model such as JC69 (Jukes and Cantor, 1969), HKY85 (Hasegawa et al., 1985) etc. Representing all the unknowns as the multidimensional parameter *θ*, this model defines probabilities *p*_*j*_ = *p*_*j*_(*θ*) for every possible pattern *j*; it is these *p*_*j*_ that are usually calculated using Felsenstein’s pruning algorithm (Felsenstein, 1973, 1981). Note in particular that *p*_*j*_ is defined for all possible *j*, including the unobserved constant patterns and any patterns that happen not to have occurred in a given data set. It is useful to write *c* = *c*(*θ*) for the total probability of occurrence of a constant site, i.e. *c* = Σ_*j*∈*c*_ *p*_*j*_, where *C* is the set containing the four constant patterns constant-A, -C, -G and -T.

The truncated data likelihood function *L*_*T*_(*θ*) is

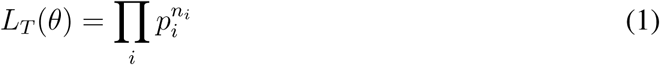

Maximizing *L*_*T*_(*θ*) over the model parameters *θ* gives their maximum likelihood (ML) estimates, 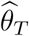, based on the truncated data. However, as shown by Lewis (2001) and Leaché et al. (2015), for SNP data sets the omission of the constant characters can cause serious estimation biases.

The problem of estimating *θ* (and *N*) in these circumstances was described by Sanathanan (1972). Following that paper, we consider the *complete* likelihood including the contribution of the *m* omitted constant sites:

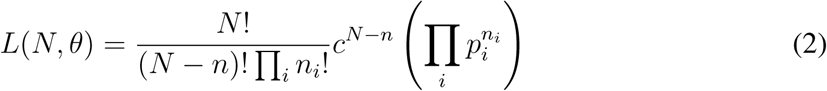

or, equivalently,

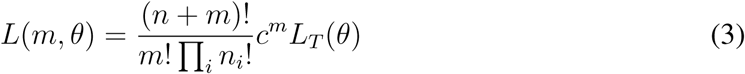

Note that this retains the combinatorial component (*n* + *m*)!/*m*! ∏_*i*_ *n*_*i*_!. In typical ML problems, where all the data are observed, the corresponding term can be omitted as it is a constant and plays no part in the maximization over *θ* (Edwards, 1972) — hence its omission from eqn. 1. However, in the truncated data case this is not true: different (inferred) values of *m* will cause the term to vary and its contribution to the likelihood cannot be ignored. Although it is unusual to infer the amount of (unobserved) data using ML, there is no reason why we should not be able to do so.

Notice that the likelihood can also be written as *L*(*m*, *θ*) = *L*_1_(*m*, *θ*)*L*_2_(*θ*) where

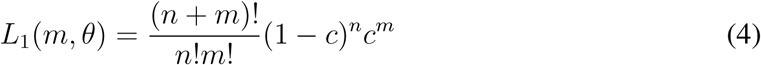

and

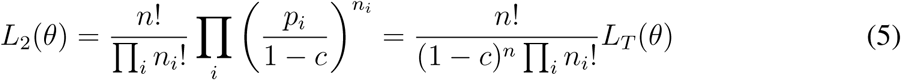

*L*_1_ is the likelihood based on the probability of *n*, and *L*_2_ is the likelihood based on the conditional probability of the *n*_*i*_ given *n* (Sanathanan, 1972).

### Conditional ML

Sanathanan (1972) describes two approaches to estimating *m* and *θ* (see also Blumenthal, 1981). The first is the method of *conditional ML* (CML), in which the conditional likelihood *L*_2_(*θ*) (eqn. 5) is maximized over *θ* to find ML model parameter estimates 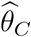. (Note that the combinatorial term *n*! /∏_*i*_ *n*_*i*_! is constant and does not affect the inference.) This corresponds to precisely the method of Felsenstein (1992) and Lewis (2001), and is equivalent to maximizing log *L*_*C*_(*θ*) over *θ*, where

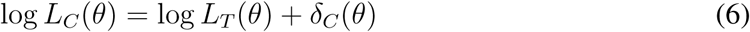

and *δ*_*C*_(*θ*) = –*n* log(1 – *c*(*θ*)) is the log-likelihood ‘correction’ term that changes the truncated data set problem into the CML problem.

In the phylogenetics context we may only be interested in the inferred tree and associated parameters 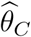. If required, however, an estimate of the number of unobserved constant sites comes from maximizing *L*_1_(*m*,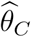) over *m* to find 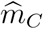. Sanathanan (1972) shows that 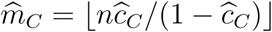 where 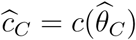 is the CML estimator of *c* and ⌊.⌋ indicates the floor function (i.e. ⌊*x*⌋ is the greatest integer ≤ *x*).

The CML version of the SNP data set phylogeny problem is implemented in RAxML v.8 (Stamatakis, 2014), invoked using the—asc-corr=lewis option (Leaché et al., 2015).

### Unconditional ML

The second approach described by Sanathanan (1972) is *unconditional ML* (UML), in which the full likelihood *L*(*m*, *θ*) (eqn. 3) is maximized simultaneously over both *m* and *θ*. We denote the corresponding inferred values by 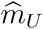 and 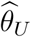.

For any fixed value *θ*\*, optimization of *L*(*m*, *θ*\*) over *m* (eqn. 3) is analogous to optimizing *L*_1_(*m*, *θ*\*) over *m* (eqn. 4) and is achieved when 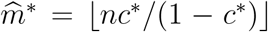 (Sanathanan, 1972). Substituting *m*\*(*θ*) = ⌊*nc*(*θ*)/(1 – *c*(*θ*))⌋ into eqn. 3 and recalling that the *n*_*i*_ are constant means the UML problem becomes one of maximizing log *L*_*U*_(*θ*) over *θ*, where

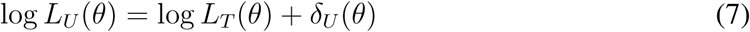

and *δ*_*U*_(*θ*) = log(*n* + *m*\*(*θ*))! – log *m*\*(*θ*)! + *m*\*(*θ*) log *c*(*θ*) is the correction term that changes the truncated problem into the UML problem.

Sanathanan (1972, 1977) proves that 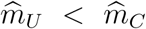 and 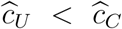, and that the asymptotic distributions of 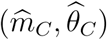 and 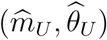 are the same. In other words, as the amount of data collected increases, the CML and UML estimators will give arbitrarily close estimates of the numbers of unobserved constant sites and model parameters. However, the approaches do not necessarily lead to equally good estimates given *finite* amounts of data.

## Results and Discussion

CML and UML both perform well: We implemented the CML and UML methods for the SNP data set phylogeny problem in order to compare their performance. We modified the baseml software (Yang, 2007) so that for each candidate value of *θ* we first compute *c*(*θ*) and *m*\*(*θ*) and then use these with the truncated log-likelihood function log *L*_*T*_(*θ*) to compute log *L*_*C*_(*θ*) and log *L*_*U*_ (*θ*) as in eqns. 6 and 7.

We simulated sequence data on the 10-taxon topology studied by Leaché et al. (2015). To create a range of realistic simulation scenarios, we scaled the tree to various lengths (scaling factors of 0.25, 0.5, 1, 2 and 4, giving tree lengths of 0.08, 0.16, 0.31, 0.62 and 1.24, respectively) and used a variety of alignment sizes (*N* = 500, 1000, 2500, 5000) under the JC69 model. After simulation, all constant site patterns were discarded. The probability of occurrence of constant sites ranged from 93% to 30% (*c* = 0.93, 0.86, 0.74, 0.54, 0.30 for scaling factors 0.25, 0.5, 1, 2, 4, respectively). For more extreme cases, inspired by population resequencing studies, we also considered scaling factors 0.03-0.21, giving rise to tree lengths 0.009-0.065 and *c* from 0.99-0.94, with *N* = 100000.

Lower scale factors lead to smaller trees and thus result in fewer variable (SNP) sites on which to base inference. Our simulations cover a range, from unobserved constant sites being rare (e.g. distantly related species, or sequencing strategies such as RADseq giving strong enrichment for variable sites) to common (e.g. closely related organisms). Our most extreme scenario (*c* = 0.99) resembles what might be observed with SNP data sets from population sequencing studies.

We used the CML and UML correction methods to re-estimate model parameters using only the variable sites from the simulated datasets, assuming knowledge of the true topology. We repeated this procedure with data simulated under the HKY85 model with moderate transition/transversion rate and nucleotide bias (*κ* = 2, *π*_*T*_ = 0.1, *π*_*C*_ = 0.2, *π*_*A*_ = 0.3, *π*_*G*_ = 0.4). In all cases, we found that estimates of branch lengths, tree length and other model parameters were almost identical with CML and UML. Figs. 1 and 2 illustrate this with a variety of results summarized from 100 simulations in each scenario; other results (not shown) all confirm these findings.

**Figure 1:**
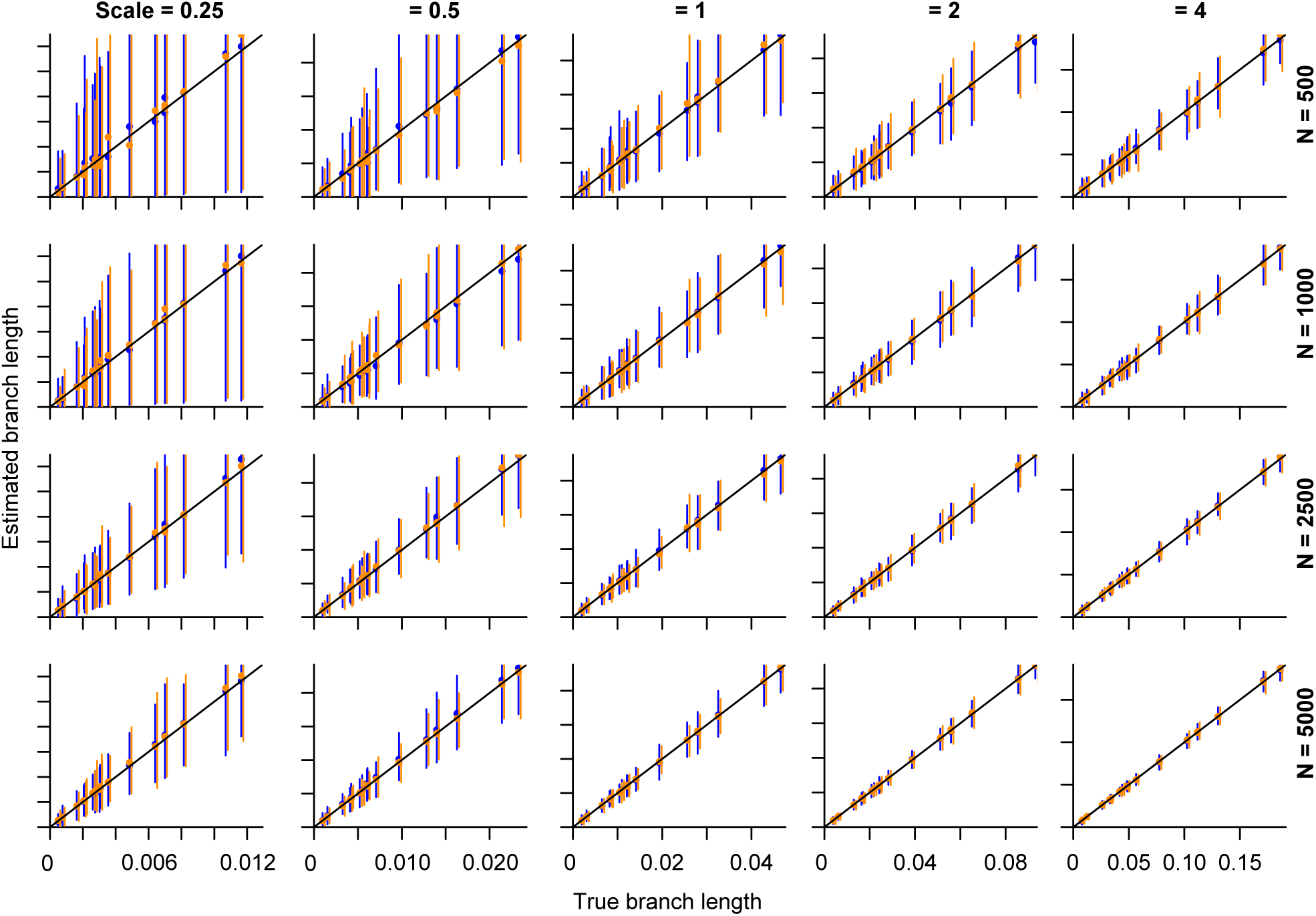
Branch length estimates under various JC69 simulation scenarios. Within each row, scenarios have the same number of sites simulated (*N*, i.e. before constant sites were removed); within columns, the same tree length scaling factor. Graphs show the mean and 5–95%-ile range for each of the 17 branch length estimates plotted against the true value, derived from 100 simulations. CML results are shown in blue; UML in orange. The two methods’ results are essentially indistinguishable.

**Figure 2:**
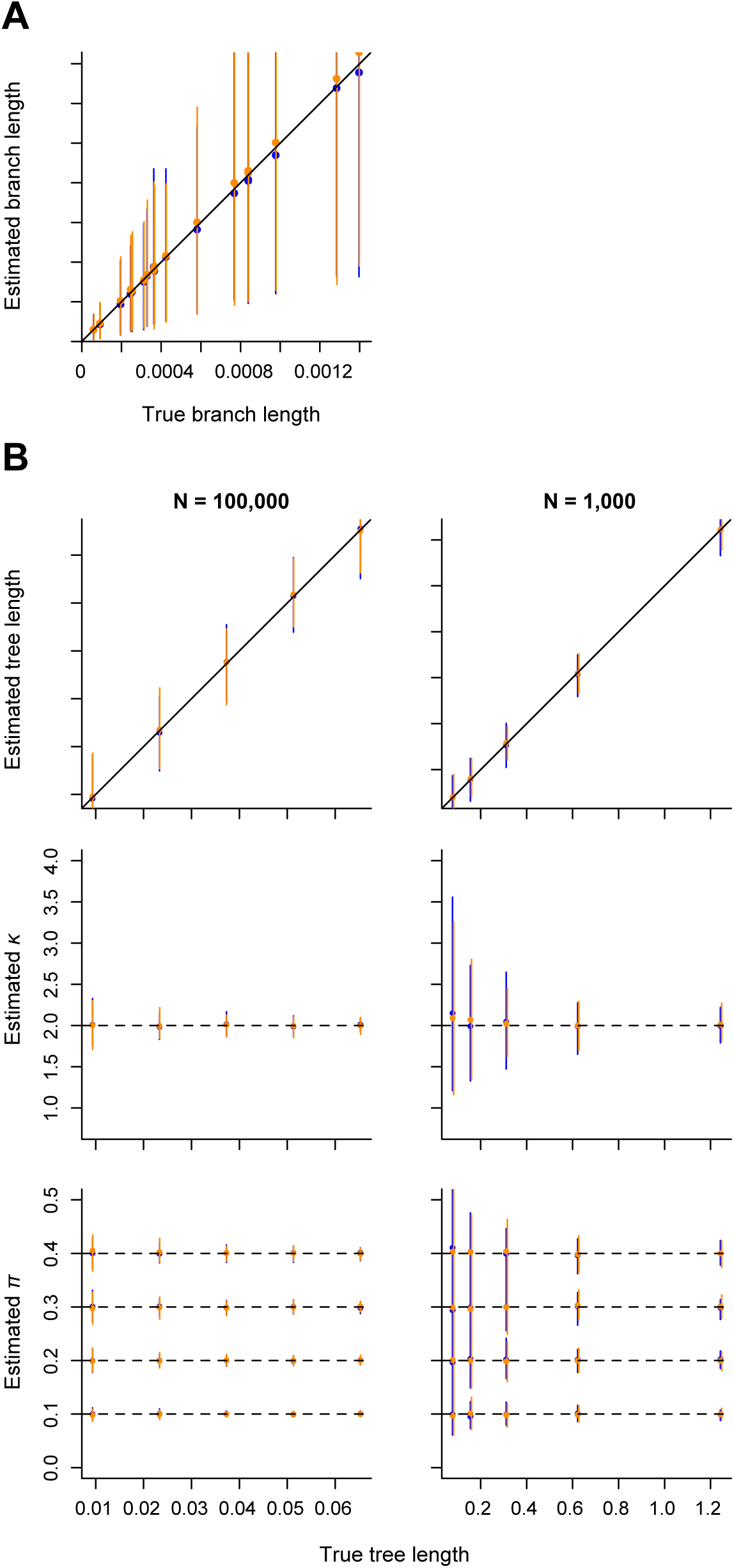
Parameter estimates (means and 5–95%-ile ranges from 100 simulations in each case) under various simulation scenarios. **A:** Branch length estimates from JC69 simulations with scaling factor 0.03 (tree length 0.09; *c* = 0.99) and *N* = 100000. **B:** Parameter estimates from HKY85 simulations. Left column: *N* = 100000, *c* from 0.99-0.94; right column: *N* = 1000, *c* from 0.93-0.30. Graphs show estimates of overall tree length (top), *κ* (middle) and nucleotide frequencies (bottom) for various tree length scaling scenarios. Colours etc. as for Fig. 1. Again, CML and UML give essentially the same results.

### Why are CML and UML almost the same?

Clearly, this is because in every case studied, the maximal values of log *L*_*C*_ (*θ*) and log *L*_*U*_ (*θ*) are attained at very similar values of *θ*. As these likelihoods differ only in the terms *δ*_*C*_(*θ*) and *δ_U_(θ*), we would like to study how these vary with *θ*. However, as this represents a complex multidimensional parameter (Yang et al., 1995), it is difficult to visualize likelihoods as *θ* varies over candidate solutions. To simplify our investigation, we give an illustration using a single JC69 simulation with the 10-taxon tree of Leaché et al. (2015) with scaling factor 1 and *N* = 1000 alignment sites simulated before removal of constant patterns. For these data, we found the truncated likelihood-optimal branch lengths 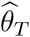 and the CML-optimal branch lengths 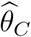, and focus attention on values of *θ* formed by interpolating between 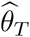 and 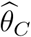 and extrapolating this range beyond 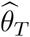 and 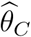. This gives a one-dimensional subspace which includes both 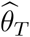 and 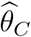. Fig. 3 shows values of log *L*_*T*_(*θ*) (eqn. 1), log *L*_*C*_(*θ*), *δ*_*C*_(*θ*) (eqn. 6), log *L*_*U*_(*θ*) and *δ*_*U*_(*θ*) (eqn. 7) for these values of *θ*, with the x-axis simultaneously labelled by the corresponding values of *c*(*θ*) and *m*\*(*θ*).

**Figure 3:**
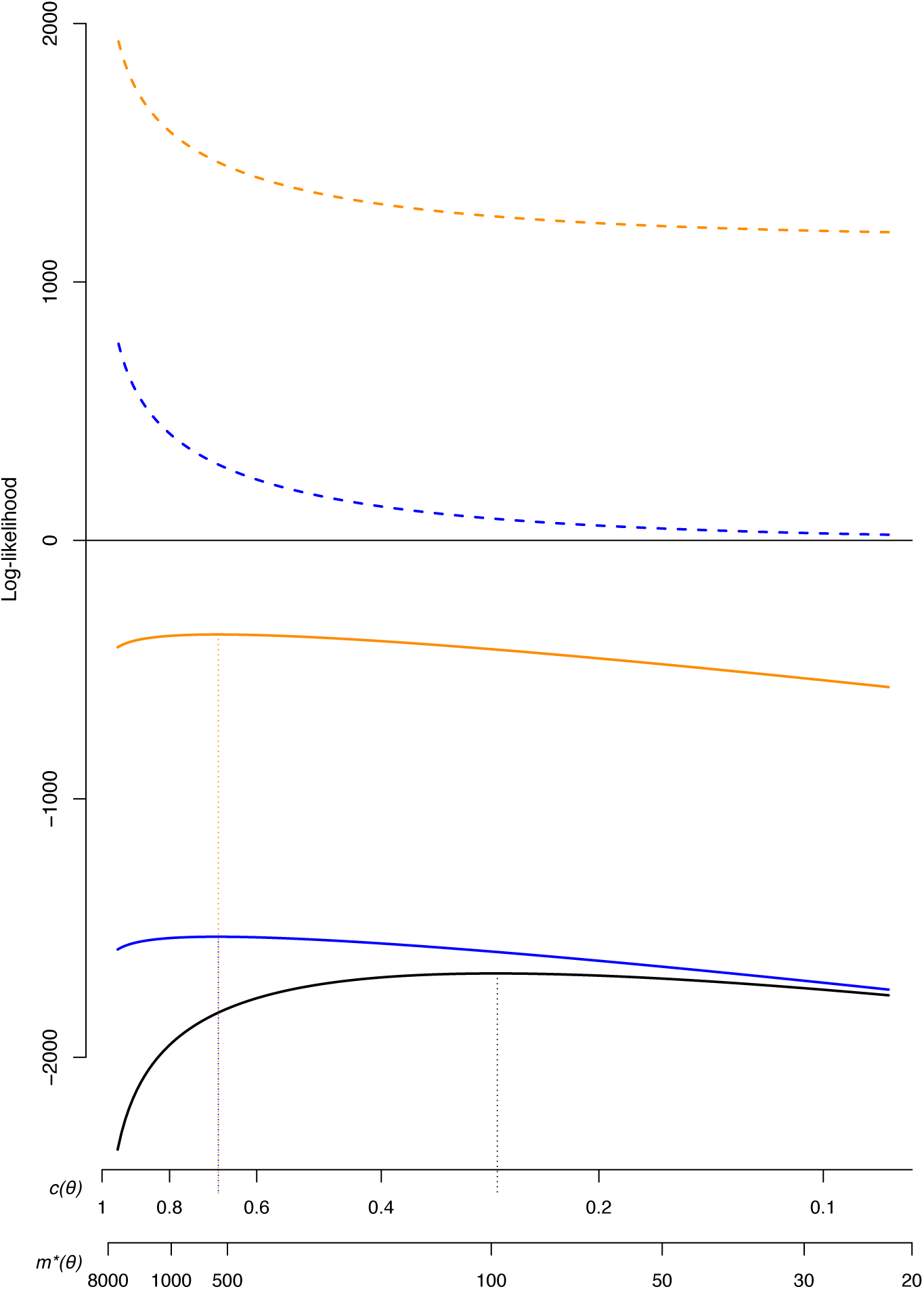
Likelihoods and correction terms in a single simulation case. From bottom to top, plots show *L*_*T*_(*θ*) (solid black line), *L*_*C*_(*θ*) (solid blue), *L*_*U*_(*θ*) (solid orange), *δ*_*C*_(*θ*) (dashed blue) and *δ*_*U*_(*θ*) (dashed orange). The *x*-axis is labelled with the values of *c*(*θ*) and *m*\*(*θ*) corresponding to the range of branch length parameters *θ* described in the text. Locations of 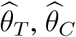 and 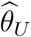 are indicated by vertical dotted lines.

Firstly, note the truncated likelihood (indicated by the solid black line) is maximized at a value of *θ* suggesting far too few omitted constant sites 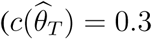 and 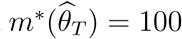, whereas the true values of *c* and *m*\* for this simulation are 0.743 and 743, respectively). This corresponds to a tree that is too divergent. The correction terms *δ*_*C*_(*θ*) and *δ*_*U*_(*θ*) (dashed blue and orange lines, respectively), while very different in absolute value, have very similar gradients. Consequently, when they are added to log *L*_*T*_(*θ*) to form log *L*_*C*_(*θ*) and log *L*_*U*_(*θ*) (solid blue and orange lines, respectively), these likelihoods have maxima at very similar values of *θ* (with 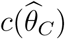 and 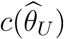 equal to 0.69, and 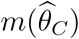 and 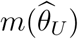 ≈ 570). In brief, the effects of the corrections *δ*_*C*_(*θ*) and *δ*_*U*_(*θ*) in the optimization of log *L*_*C*_(*θ*) and log *L*_*U*_(*θ*) are indeed very similar.

It is not the absolute values of *δ*_*C*_ and *δ*_*U*_ that are critical to the difference between 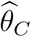 and 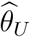, but how they vary in the regions of the CML and UML optima. Further analysis might consider the derivatives of *δ*_*C*_ and *δ*_*U*_ with respect to *θ*, but in the SNP phylogeny question this is complicated by the non-standard form of the topology parameter *θ* (Yang et al., 1995) and by the factorial and floor functions in *δ*_*U*_. However, *δ*_*C*_ and *δ*_*U*_ only depend on *θ* through the probability of constant patterns *c*(*θ*) and so an alternative approach is to consider the relative variation of *δ*_*C*_ and *δ*_*U*_ as *c* varies. In particular, if *δ*_*C*_(*c*_1_) – *δ*_*C*_(*c*_0_) and *δ*_*U*_(*c*_1_) – *δ*_*U*_(*c*_0_) are very similar for any *c*_0_ ≈ *c*_1_ (and in particular near to 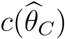 and 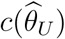), then the CML and UML correction terms will have similar effects on the optimizations of log *L*_*C*_ and log *L*_*U*_, leading to similar CML and UML estimates.

Indeed, it can be shown (omitted for brevity) that if *c*_0_ and *c*_1_ are chosen to be similar, such that the corresponding values 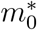 and 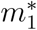 satisfy 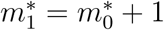, then

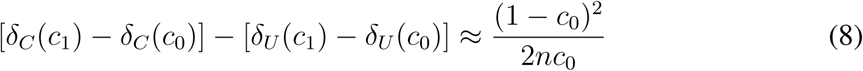

This represents a measure of the difference between the gradients *dδ*_*C*_/*dm* and *dδ*_*U*_/*dm* (Fig. 3); since it scales as 1/*n*, *δ*_*C*_(*θ*) and *δ*_*U*_(*θ*) will be expected to have very similar effects for reasonably large values of *n*, which will be the case for most phylogenetic problems.

### Base frequencies can behave unexpectedly in SNP datasets

While analyzing simulated datasets as described above, we noticed the observed frequencies of nucleotides A, C, G and T in the HKY simulations did not always match the corresponding model parameters. We realized this is because in those models where different nucleotides have different substitution rates, the constant-A, -C, -G and-T site patterns have different probabilities of occurrence. As a consequence, the observed frequencies of A, C, G and T amongst the constant site patterns omitted from SNP datasets, and amongst the SNP patterns retained for analysis, will differ from the model parameter values. This is illustrated in Fig. 4, using the HKY simulation scenario described above.

**Figure 4:**
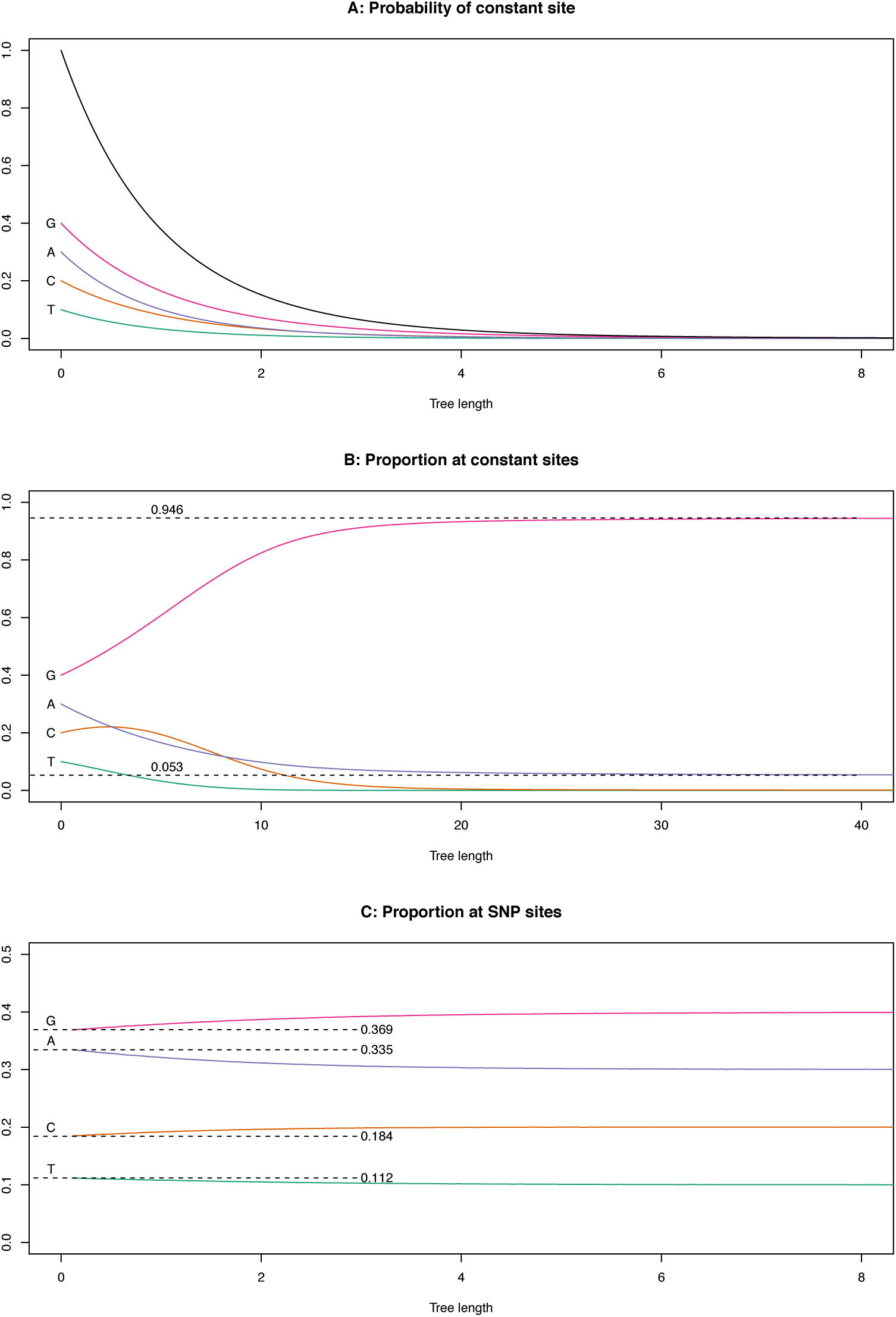
Base frequencies observed with SNP datasets. **A:** As the 10-taxon tree used throughout this study is scaled from a length of 0 to increasingly large branch lengths, the proportion of constant sites (top, black) falls to 0; the component proportions of constant-A,-C,-G and-T patterns falls from their equilibrium values to 0. **B:** Conditional on observing a constant pattern, the proportions of nucleotides A, C, G and T vary as the tree size increases. (Note that these proportions need not vary monotonically, as observed for the constant-C patterns in this example.) **C:** Conditional on observing a non-constant (SNP) pattern, the observed base frequencies also differ from the model parameters for shorter tree lengths.

As a consequence, it may be advisable to estimate base frequencies using ML rather than the simple counting (empirical) method when working with SNP data (Goldman, 1993). We have used this approach throughout this paper.

## Conclusions

In all of our analyses of simulated data using the CML and UML approaches for removing ascertainment bias from SNP datasets, we have found virtually no difference in the results obtained. We explain this by observing that the different approaches’ respective correction terms *δ*_*C*_ and *δ*_*U*_ behave very similarly in their effects (Fig. 3). This has further been supported by our analysis of the gradients of *δ*_*C*_ and *δ*_*U*_, which confirms their similarity for plausible phylogenetic scenarios and data quantities. Although we have not investigated tree topology estimation, the near-identical results of CML and UML for estimation of other parameters, including branch lengths, lead us to think it very unlikely that they could behave differently for topology estimation. We therefore conclude that it is of little importance which of these methods is used in practice in phylogenetic studies. The CML method is widely available via the—asc-corr=lewis option of RAxML (Stamatakis, 2014; Leaché et al., 2015).

We note in passing that the observed base frequencies in SNP datasets can differ systematically from the corresponding substitution model parameters, due to bias in the frequency with which constant-nucleotide site patterns arise and are thus omitted. A simple solution to this should be to use ML to estimate these parameters (Goldman, 1993).

## Acknowledgments

We thank Joe Felsenstein and Alexis Stamatakis for discussions of an earlier (incorrect) attempt at this study, Jakub Truszkowski for helpful discussions of that attempt and then the CML and UML methods, and Melissa Ward for alerting us to issues of base frequency estimation with SNP datasets.

